# Homeostatic plasticity triggered by rod photoreceptor degenerative disease is associated with maintenance of sensitive night vision

**DOI:** 10.1101/2020.05.28.121673

**Authors:** Henri Leinonen, Nguyen C Pham, Taylor Boyd, Johanes Santoso, Krzysztof Palczewski, Frans Vinberg

**Author notes:** Equal contribution (Henri Leinonen, Nguyen C Pham). **To whom correspondence should be addressed**: Frans Vinberg, Ph.D., John A. Moran Eye Center, Dept. of Ophthalmology & Visual Sciences, University of Utah, 65 Mario Capecchi Dr., UT 84132, Henri Leinonen, Ph.D., Gavin Herbert Eye Institute, Dept. of Ophthalmology, University of California, Irvine, CA 92657.

## Abstract

Neuronal plasticity of the inner retina has been observed in response to photoreceptor degeneration. Typically, this phenomenon has been considered maladaptive and may preclude vision restoration in the blind. However, several recent studies utilizing triggered photoreceptor ablation have shown adaptive responses in bipolar cell dendrites expected to support normal vision. Whether such homeostatic plasticity occurs during progressive photoreceptor degenerative disease to help maintain normal visual behavior is unknown. We addressed these issues in an established mouse model of Retinitis Pigmentosa caused by the P23H mutation in rhodopsin. We show robust modulation of the retinal transcriptomic network reminiscent of the neurodevelopmental state as well as potentiation of rod – rod bipolar cell signaling following rod photoreceptor degeneration. Additionally, we found highly sensitive night vision in P23H mice even when more than half of the rod photoreceptors were lost. The results implicate retinal adaptation leading to persistent visual function during photoreceptor degenerative disease.

## INTRODUCTION

Neurons and neuronal networks require mechanisms for maintaining their stability in the face of numerous perturbations occurring over a lifetime. Homeostatic plasticity refers to the process whereby the activity of the neuron or neuronal network is maintained (Burrone & Murthy, 2003; G. Turrigiano, 2012). Homeostatic plasticity is a relatively new concept as it was first described in the central nervous system in 1998 (G. G. Turrigiano, Leslie, Desai, Rutherford, & Nelson, 1998). Since then various homeostatic plasticity mechanisms have been reported in different regions of the brain, including the hippocampus and somatosensory and visual cortices, using *in vivo* models and *ex vivo* and *in vitro* preparations (Gainey & Feldman, 2017; G. Turrigiano, 2012; G. G. Turrigiano, 2017). As examples, binocular visual deprivation triggers compensatory synaptic changes in the primary visual cortex leading to the stabilization of spiking activity (Desai, Cudmore, Nelson, & Turrigiano, 2002; Goel et al., 2006; Goel & Lee, 2007), and stroke can trigger homeostatic plasticity that is thought to compensate for neuronal damage, which plays an important role in early phase rehabilitation (Murphy & Corbett, 2009). Homeostatic plasticity counteracts insufficient activity in neural synapses (Burrone & Murthy, 2003; G. Turrigiano, 2012), but it also works to prevent saturation of synapses that could be caused by positive feedback mediated by Hebbian plasticity, which is better known for its role in experience-dependent plasticity, learning and memory (Abraham & Bear, 1996).

The neural retina of the eye is considered relatively stable both structurally and functionally after postnatal development. Still, remodeling of the retina in response to photoreceptor degenerative diseases is well established. Retinal remodeling often has been described as a maladaptive process that can exacerbate the disease and impede treatment strategies designed to rescue light responsiveness of the retina (Fariss, Li, & Milam, 2000; Jones & Marc, 2005; Marc & Jones, 2003; Telias et al., 2019; Toychiev, Ivanova, Yee, & Sagdullaev, 2013). It involves formation of ectopic synapses and expression of disruptive spontaneous oscillatory activity in the post-receptoral inner retina during photoreceptor degenerative disease (Goo et al., 2011; Haverkamp et al., 2006; Margolis, Newkirk, Euler, & Detwiler, 2008; Michalakis et al., 2013; Stasheff, 2008; Toychiev et al., 2013; Tu, Chen, McQuiston, Chiao, & Chen, 2015). In contrast, recent evidence supports adaptive rather than solely destructive neuronal changes. For example, dendritic compensatory synapse formation occurs in retinal bipolar cells when various retinal cells, including rod and cone photoreceptors, have been ablated (Beier et al., 2017; Beier, Palanker, & Sher, 2018; Care et al., 2019; Johnson et al., 2017; Shen, Wang, Soto, & Kerschensteiner, 2020). These studies suggested that rod or cone bipolar cell dendrites can extend their arbors and form new synapses with their correct targets (*i.e.* rod and cone bipolar cells) either within their normal territory or sometimes outside of their normal dendritic field to compensate for the reduced number of photoreceptor cells. These homeostatic changes could help maintain retinal output and vision at or near normal levels. It is likely that the mechanisms, onset and time course of degeneration as well as the cell types in question all contribute to remodeling in the retina, and whether it will have a supportive or deleterious effect on vision. Thus, it will be critical to determine the nature of remodeling and plasticity across various retinal degenerative conditions.

Whether homeostatic plasticity is present during a progressive retinal degenerative disease is unknown. We hypothesized that an inherited photoreceptor degenerative disease that progresses in a moderate manner would promote homeostatic plasticity supporting normal vision. We tested this hypothesis using an established mouse model of the most common form of Retinitis Pigmentosa (RP) caused by a dominant P23H mutation in the rod visual pigment, rhodopsin (Sakami, Kolesnikov, Kefalov, & Palczewski, 2014; Sakami et al., 2011). By conducting whole retinal RNA-sequencing, *in vivo* and *ex vivo* electrophysiology, as well as behavioral experiments in diseased and control mice during the course of disease progression, we demonstrate that rod photoreceptor degeneration triggers homeostatic plasticity and the potentiation of rod–rod bipolar cell signaling that is associated with the maintenance of highly sensitive scotopic vision. This type of compensatory mechanism may explain why patients with inherited retinal diseases can maintain normal vision until a relatively advanced disease state is reached and could inspire novel treatment strategies for blinding diseases.

## RESULTS

### Time course of rod degeneration and functional decline in heterozygous P23H mice

Retinal remodeling following photoreceptor cell death has been observed in a number of genetic and induced animal models of retinal degeneration (Beltran, 2009; Chang, Hawes, Davisson, & Heckenlively, 2007; LaVail et al., 2018; Petersen-Jones, 1998; Ross et al., 2012), as well as in *post mortem* human eye specimens from patients with Retinitis Pigmentosa (RP) or Age-related Macular Degeneration (AMD) (Fariss et al., 2000; Jones, Pfeiffer, Ferrell, Watt, Marmor, et al., 2016; Jones, Pfeiffer, Ferrell, Watt, Tucker, et al., 2016; Li, Kljavin, & Milam, 1995). In the current study, we utilized a well-established mouse model of autosomal dominant RP (Sakami et al., 2014; Sakami et al., 2011), caused by a heterozygous P23H mutation in the rhodopsin gene (*Rho^P23H/WT^*). These mice will be referred to in this report as P23H mice. We started by confirming the suitability of this model for our study, focusing on rod-mediated retinal signaling and night vision. We followed structural changes of the retina by standard histology and optical coherence tomography (OCT) imaging, and confirmed that the outer nuclear layer (ONL) where the photoreceptor nuclei reside, and the inner and outer segments (IS and OS, respectively) of rod and cone photoreceptors progressively degenerated, while inner retinal layers appeared to remain anatomically intact (Figure 1A-G). In mice, the thickness of the ONL correlates with the number of surviving rod photoreceptors. We found 20 %, 60 % and 73 % decreases in central retinal ONL thickness at 1-, 3- and 5-months of age, respectively, in P23H mice compared to WT littermates (Figure 1H). Next, using *in vivo* electroretinography (ERG) we tested the light-activated mass electrical response arising from the retina. We analyzed the a-wave of the fully dark-adapted retina, which is primarily a rod-dominant response, although having a minute contribution from cone cells. The dark-adapted a-wave showed an almost 50 % suppression in P23H mice by 1-month of age (Figure 1I, J) consistent with an earlier study by Sakami et al. (Sakami et al., 2011).

**Figure 1.**
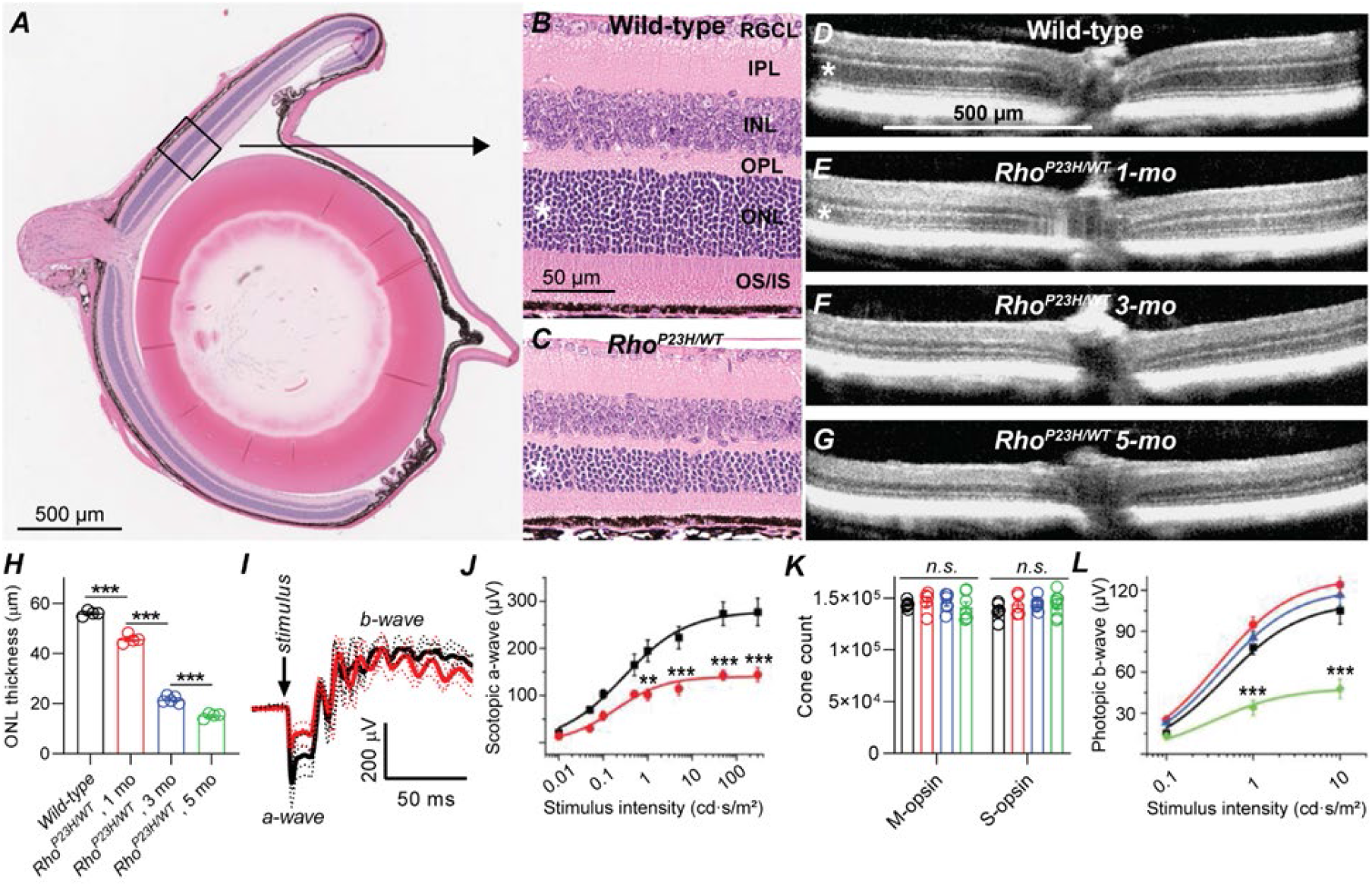
Characterization of retinal degeneration progression in heterozygote P23H rhodopsin mutated mice. (A) Representative histology image of a wild-type (WT) mouse eye. (B-C) Representative magnified histology images of 1-month-old WT (B) and P23H (C) dorsal retinas. Asterisks mark the ONL, *i.e.* photoreceptor nuclei layer. RGCL, retinal ganglion cell layer; IPL, inner plexiform layer; INL, inner nuclear layer; OPL, outer plexiform layer (the site of photoreceptor-bipolar cell synapse); IS/OS, photoreceptor inner and outer segments. (D-G) Representative optical coherence tomography images. (H) ONL thickness analysis from central retina. Data averaged over for retinal quadrants. (I) Scotopic ERG waveforms in response to 100 cd•s/m2 flash in 1-month-old WT (black) and P23H (red) mice. Data are averaged from 4 mice per group. (J) ERG a-wave amplitude analysis at 1-month of age. (K) Total M- and S-opsin counts in whole mount retinas (WT, black; P23H 1 mo, red; 3 mo, blue; 5 mo, green) (L) Photopic ERG b-wave amplitude analysis (WT, black, n=6; P23H 1 mo, red, n=8; 3 mo, blue, n=7; 5 mo, green, n=5). Bonferroni posthoc tests: \**P*<0.05, \*\**P*<0.01, \*\*\**P*<0.001.

Cone cells comprise only 3 % of the photoreceptor population in pigmented mice. Therefore, the experiments just described lacked the necessary sensitivity to evaluate cone cell survival. To investigate the fate of cone cells in P23H mice, we used an immunohistochemical approach to determine the number of cone cells and light-adapted, cone-dominant ERG recordings to assess their functional status. The total count of short- and medium-wavelength sensitive S- and M-cones, respectively, remained constant even at the latest time point tested, at 5-months of age (Fig. 1K). ANOVA analysis revealed a modest but statistically significant (F_1,12_=5.25, *P*=0.04) increase in the photopic ERG amplitude in P23H mice compared to WT littermates at 1-month of age, which continued at 3-months of age (Figure 1L). However, cone function started to decline thereafter, and at 5-months of age the photopic ERG amplitudes significantly decreased in P23H mice.

Taken together, the rod photoreceptors in the P23H mouse model of RP are preferentially targeted in the early stage of disease, and the loss of these rod photoreceptors occurs at a moderate rate. Therefore, the model is suitable for investigating the compensatory network changes that arise during progressive rod cell loss.

### RNA-sequencing from retinas reveals a robust neuronal network adaptation in early RP

We initially used RNA-sequencing (RNA-seq) to determine whether the transcriptomic profile of the retina changed in a manner that might provide functional adaptation in response to rod photoreceptor degeneration. We performed RNA-seq at 1-month since at this age the P23H mouse retina has matured but has not yet degenerated severely (Figure 1H). We found 2721 downregulated and 2683 upregulated genes in P23H mice compared to WT littermates (Figure 2A). There were few differentially expressed genes between sexes, unrelated to disease, and therefore we pooled samples from both genders in the remainder of the analyses.

**Figure 2.**
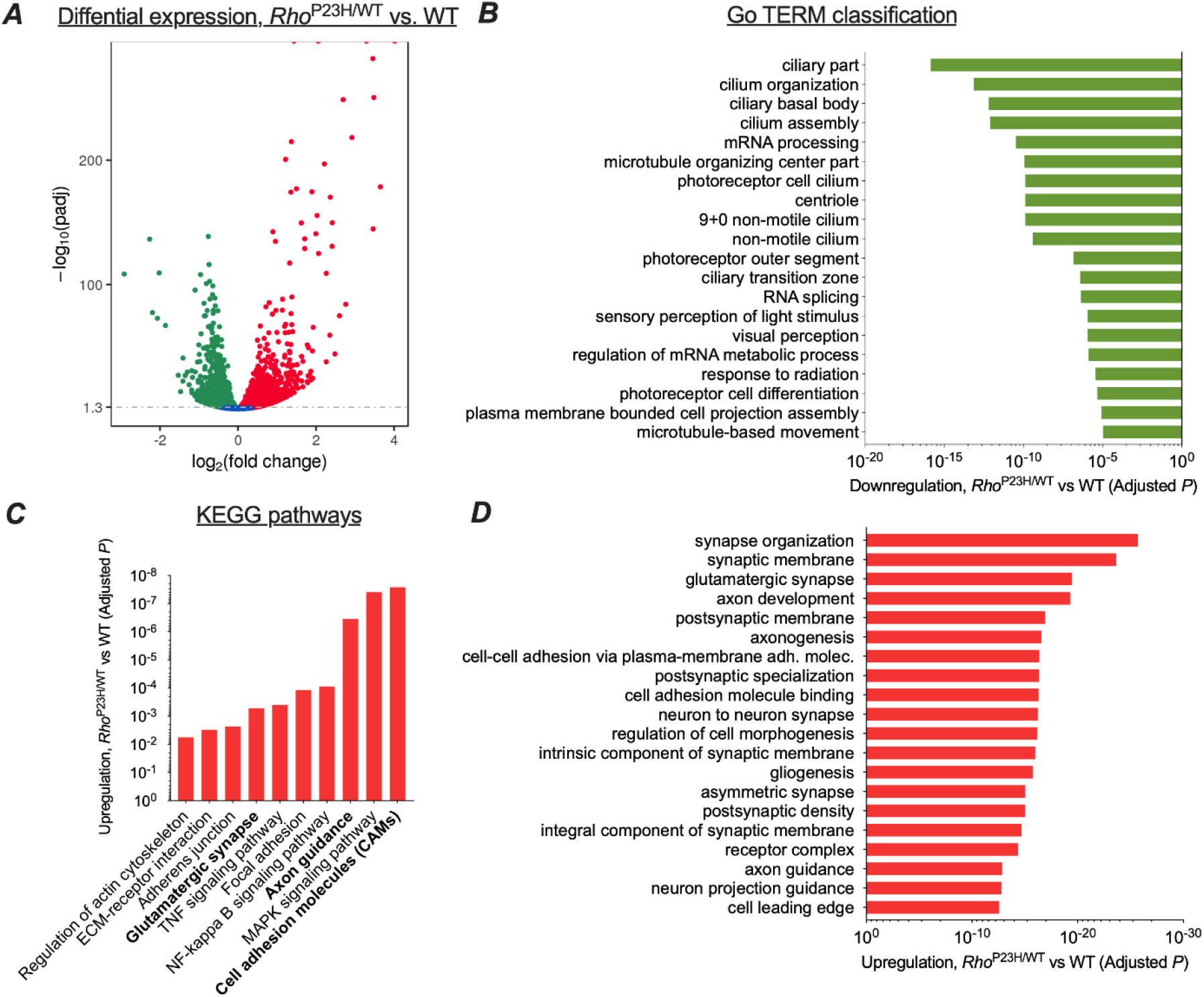
Transcriptomic analysis shows neural network adaptation to rod death in early stage retinitis pigmentosa. RNA-sequencing was performed from whole retinal extracts at 1-month of age in P23H mice (n=7) and WT (n=7) littermates. (A) A total of 5404 transcripts showed differential expression (downregulation, green; upregulation, red) between P23H and WT retinas. (B) Gene ontology (GO) term analysis shows expected downregulation in photoreceptor-dominant gene clusters. (C) KEGG analysis shows significant enrichments in cell/neuron adhesion and growth pathways, glutamatergic synapse formation and several pathways associated with both cell stress and synaptic plasticity. (D) GO term analysis illustrates the most significant upregulation in synaptic and neuronal development and growth pathways in P23H retinas.

We used gene ontology analysis (GO) to objectively identify the 20 most downregulated and 20 most upregulated gene classes in P23H mouse retinas based on the *P*-values. As expected, numerous photoreceptor-specific gene classes were significantly downregulated in P23H mice, such as those related to ciliary structure, phototransduction, and photoreceptor cell differentiation (Figure 2B). In contrast, GO analysis revealed robust increases in gene clusters associated with neuronal growth and development, encompassing such cellular processes as synapse organization, postsynaptic specialization, glutamatergic synapse formation, axonogenesis, and cell-cell adhesion (Figure 2D). Furthermore, KEGG pathway analysis showed a total of 74 statistically upregulated pathways including those involved in cell adhesion, axon guidance, and glutamatergic synapse processing (Figure 2C), consistent with neuronal network adaptation following the initial sensory defect in the retina. As expected, several pathways associated with cell stress were also upregulated, including the MAPK, NF-kappa B, and TNF signaling pathways. To verify the activation of at least one of these key pathways, the MAPK signaling pathway, ERK phosphorylation was analyzed in whole retina extracts and found to be enhanced in P23H mice (Figure 2 – figure supplement 1).

### Increased sensitivity of rod bipolar cells to their rod input in P23H mice

To evaluate if transcriptomic network adaptation in P23H mice in response to rod degeneration is associated with changes in retinal light signaling, we first recorded light-evoked responses from isolated mouse retinas using *ex vivo* ERG (Figure 3A). E*x vivo* ERG allows quantitative dissection of photoreceptor (*R_PR_*) and ON bipolar cell (*R_BC_*) responses by using blockers for the metabotropic glutamate receptor (mGluR) and potassium channels in Müller glia cells (Bolnick, Walter, & Sillman, 1979; Vinberg, Kolesnikov, & Kefalov, 2014), see Materials and Methods, Figure 3B-D).

**Figure 3.**
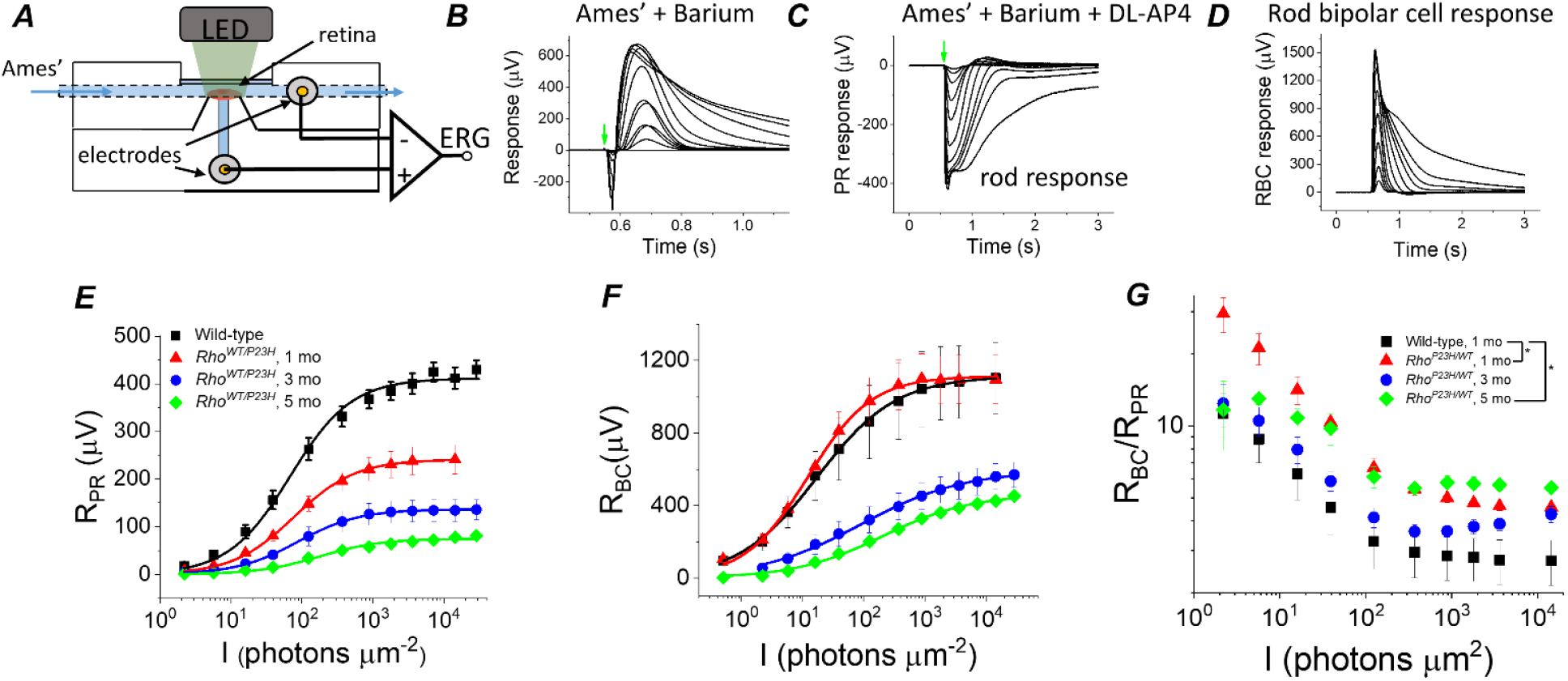
Sensitivity of bipolar cell responses to their photoreceptor input increases in P23H mice. (A) Custom *ex vivo* ERG specimen holder used to record transretinal voltage with two Ag/AgCl macro electrodes. Isolated retina without retinal pigment epithelium is mounted photoreceptor side up and perfused with Ames’ medium. (B) Representative light flash responses from a dark-adapted WT control mouse retina containing 100 μM BaCl_2_ to remove signal component arising from Müller cells. (C) Light responses from the same retina after addition of DL-AP4 eliminating metabotropic glutamatergic transmission to reveal the photoreceptor-specific responses. (D) Bipolar cell responses obtained by subtracting the traces shown in (C) from those in (B). Amplitude data for photoreceptor (R_PR_, E) and bipolar (R_BC_, F) responses, and R_BC_/R_PR_ ratio (G) as a function of light flash intensity in photons μm^-2^ (505 nm). Mean ± SEM. In panel G, \**P*<0.05 (two-way ANOVA between-subjects effect). Smooth traces in E and F plot Eq (1) fitted to the mean amplitude data. Statistics for *R_max_* and intensity required to generate half-maximal photoreceptor or ON bipolar cell response (*I_I/2_*) are in Table 1. (Control, 1 mo, n=4; P23H, 1 mo, n=4; 3 mo, n=3; 5 mo, n=4 mice).

As expected, we found that the maximal photoreceptor response decreased by ~40 % at 1-month of age in P23H mice as compared to control mice, and responses continued to decrease thereafter (Figure 3E, Table 1). These changes, initially at 1-month of age, were larger than expected when compared to the total loss of rod photoreceptors (Figure 1H), suggesting that the decrease in the light sensitive current of the rods precedes rod degeneration in this model. Interestingly; however, light sensitivity measured as the light intensity to generate a half-maximal response (*I_1/2_*) was not reduced in 1-3-month-old P23H mice (Figure 3E, Table 1). At 5 months of age, *I_1/2_* of P23H mice significantly increased (Figure 3E, Table 1). This may be due to the contribution from cone cells at this age (Figure 1I and K). Progressive reduction of rod photoreceptor light responses was expected and has been reported previously (Sakami et al., 2011). However, in this study the main issue was to determine whether the observed transcriptomic changes correlate with changes in how bipolar cells respond to their photoreceptor input in P23H mice. Strikingly, despite the ~40% decrease of photoreceptor input in 1-month-old P23H mice, their bipolar cell responses did not diminish at any of the light flash intensities utilized (Figure 3F). Since rod bipolar cells derive their responses by pooling input from several rod photoreceptors, we quantified the sensitivity of rod bipolar cells to their intrinsic input in two ways. First, we compared the ratio of bipolar cell and photoreceptor cell responses (*R_BC_*/*R_PR_*) between control and P23H mice at each light flash intensity (Figure 3G). This ratio increased in 1-month old P23H mice compared to control mice, indicating a significant increase in the sensitivity of the bipolar cells. We also found less pronounced but statistically significant increases of *R_BC_*/*R_PR_* in 5-month-old P23H mice (Figure 3G). These results are consistent with two prior studies showing less severe reduction of the *in vivo* ERG b-waves as compared to a-waves in a P23H transgenic rat model (Aleman et al., 2001; Machida et al., 2000). However, one caveat in analyzing *R_BC_*/*R_PR_* is that the photoreceptor response amplitudes are not linear functions of light intensity. Thus, a second, more quantitative way to determine the sensitivity of bipolar cells to their photoreceptor input is to plot *R_BC_* as a function of *R_PR_* and determine the *R_PR_* required to generate a half-maximal *R_BC_* (=*R_P,1/2_*, see Figure 4D-F below and Materials and Methods). This analysis revealed a significant decrease of *R_P,1/2_*, *i.e.* sensitization of bipolar cells in P23H mice at all ages tested (1, 3 and 5 months) as compared to WT control mice (Table 1).

**Figure 4.**
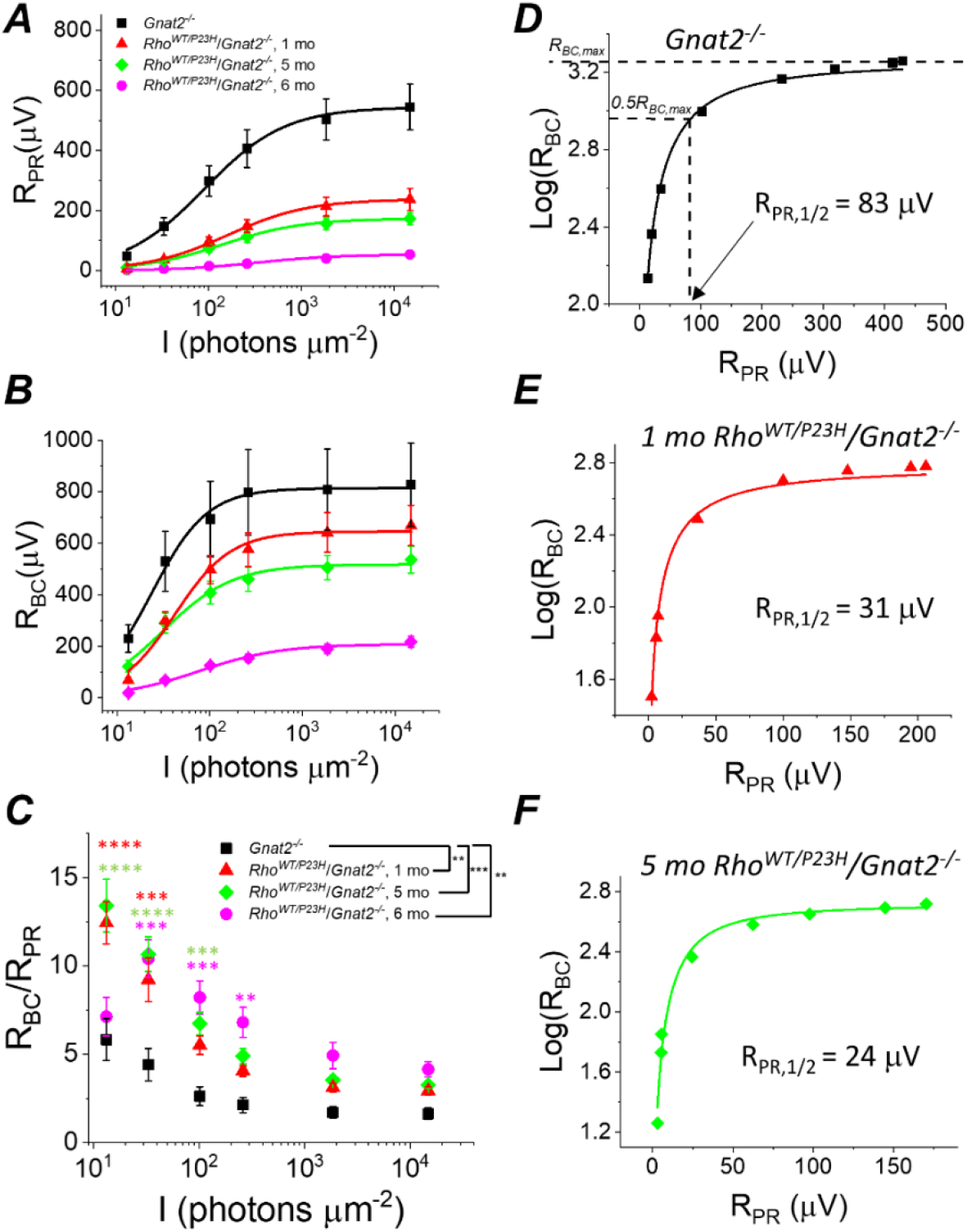
Sensitivity of rod bipolar cell responses to their rod input increases in P23H/*Gnat2^-/-^* mice. *Ex vivo* ERG rod (A) and RBC (B) response amplitudes, and their ratio (C) derived as in Fig. 3. (D-F) RBC plotted as a function of R_PR_ in individual retinas from *Gnat2^-/-^* control (D), and 1 mo (E) and 5 mo (F) P23H/*Gnat2^-/-^* mice. Smooth lines in A and B plot Eq. (1) fitted to mean data. Smooth lines in D-F plot Eq. (3) fitted to data from individual retinas. \**P*<0.05, \*\**P*<0.005, \*\*\**P*<0.001, \*\*\*\**P*<0.0001. Statistics for *R_max_* and intensity required to generate half-maximal rod or RBC response (*I_1/2_*) as well as *R_PR,1/2_* are in Table 1. In C, two-way ANOVA followed by Bonferroni’s posthoc test was used to compare between-subjects main effect and the intensity where a significant effect was found, respectively. (Pooled control, 1-6 mo, n=7(11); P23H, 1 mo, n=4(6); 5 mo, n=11(18); 6 mo, n=2(4) mice(retinas)).

**Table 1.** Ex vivo ERG parameters. *I_1/2_,* photons (505 nm) per flash/μm^2^ from fitting Eq. (1) to photoreceptor (pharmacologically isolated) and bipolar cell (subtracted) amplitude data; *R_max_,* maximum amplitude measured at plateau of the photoreceptor response and at the peak of the bipolar cell component in response to bright flash (~15,000 photons μm^-2^); *R_PR,1/2_*, photoreceptor response amplitude required for half-maximal bipolar cell response as determined by fitting Eq. (3) to R_PR_ – R_BC_ amplitude data. C57 background: Control, 1 mo, n=4; P23H, 1 mo, n=4; 3 mo, n=3; 5 mo, n=4 mice; *Gnat2^-/-^* background: Pooled control, 1-6 mo, n=7(11); P23H, 1 mo, n=4(6); 5 mo, n=11(18); 6 mo, n=2(4) mice(retinas). \**P*<0.05, \*\**P*<0.005, \*\*\**P*<0.001 in comparison to control data, two-tailed t-test. All data mean ± SEM.

| Genotype/age | Photoreceptors |  | Bipolar cells |  |  |
| --- | --- | --- | --- | --- | --- |
| | $I_{1/2}$ (hv/$\mu\text{m}^2$) | $R_{max}$ ($\mu\text{V}$) | $I_{1/2}$ (hv/$\mu\text{m}^2$) | $R_{max}$ ($\mu\text{V}$) | $R_{PR,1/2}$ ($\mu\text{V}$) |
| 1 mo Control P23H/C57 | 74 ± 18 | 410 ± 20 | 24 ± 8 | 1,120 ± 203 | 76 ± 9 |
| 1 mo P23H/C57 | 78 ± 10 | 242 ± 29** | 15 ± 2 | 1,152 ± 40 | 29 ± 6** |
| 3 mo P23H/C57 | 98 ± 13 | 138 ± 24*** | 71 ± 30 | 547 ± 74* | 39 ± 1* |
| 5 mo P23H/C57 | 285 ± 83* | 81 ± 7*** | 72 ± 7* | 388 ± 14* | 24 ± 1** |
| Pooled Control P23H/Gnat2 | 104 ± 17 | 545 ± 76 | 33 ± 12 | 830 ± 160 | 68 ± 9 |
| 1 mo P23H/Gnat2 | 230 ± 40** | 238 ± 37** | 48 ± 6 | 670 ± 80 | 38 ± 6* |
| 5 mo P23H/Gnat2 | 214 ± 32* | 176 ± 21*** | 42 ± 5 | 540 ± 50 | 28 ± 4*** |
| 6 mo P23H/Gnat2 | 339 ± 47*** | 54 ± 8*** | 88 ± 9** | 220 ± 20** | 13 ± 2*** |

Since the mouse retina is rod-dominant, our results strongly suggest that rod bipolar cells become more sensitive to their rod input in P23H mice. However, at an older age, it is possible that the cone-mediated responses, which we found to be preserved longer compared to rods (Figure 1J-K), and even hyper-sensitized in the early disease state (Figure 1K), could dominate and explain the compensation of bipolar cell responses in P23H mice. To exclude this possibility and to study more quantitatively the implications of the P23H rhodopsin mutation in the rod signaling pathway, we crossed P23H mice with a cone transducin-α (*Gnat2*) knock-out (P23H/*Gnat2^-/-^*) mouse line. The cones in these mice do not generate light responses but have no structural abnormalities (Ronning et al., 2018). We observed a qualitatively similar phenomenon in P23H/*Gnat2^-/-^* mice as in P23H knock-in mice (Figures 3, 4). The maximal rod response amplitude was suppressed by more than half in 1-month-old P23H/*Gnat2^-/-^* mice compared to their littermate control mice (Figure 4A). A further decrease of the maximal rod response amplitudes in aging P23H/*Gnat2^-/-^* mice was accompanied by the desensitization of their rods as indicated by larger values of *I_1/2_* in 1 – 6-month-old P23H/*Gnat2^-/-^* mice compared to their littermate controls (Figure 4A, Table 1). The rod bipolar cell responses decreased less than the rod responses in 1- and 5-month-old P23H/*Gnat2^-/-^* mice before a relatively steep decrease from 5 to 6 months of age (Figure 4B). Since the decrease of *R_BC_* was slower compared to the decline in *R_PR_* with age, the *R_BC_/R_PR_* ratios remained significantly higher up to 6-months of age in P23H/*Gnat2^-/-^* mice (the oldest age tested, Figure 4C). Moreover, we found an almost two-fold decrease in photoreceptor input required to generate a half-maximal RBC response in 1-month-old P23H/*Gnat2^-/-^* mice as compared to that in control mice, and *R_PR,1/2_* continued to decrease down to ~20% of that in control retinas in 6-month-old P23H mice (Figure 4D-F, Table 1).

Although the cones lacking GNAT2 do not generate light responses, light signal transmission from rods to the cone pathway through gap junctional coupling could be upregulated in P23H mice to compensate for the reduced rod input. To test this hypothesis, we determined the effect of a gap junction blocker, meclofenamic acid (MFA), on the photoreceptor a-wave and ON bipolar cell b-wave by *ex vivo* ERG in P23H/*Gnat2^-/-^* mice (Figure 5A,B). If potentiation of the b-wave responses is due to increased signaling of rods to the cone pathway via gap junctions, we expected to see suppression of the b-wave amplitudes by MFA. Although MFA trended towards suppressing the b-wave amplitude, it appeared to cause even larger suppression of the a-wave amplitude (Figure 5B). This result suggests that electrical transmission from rods to cones or cone bipolar cells does not contribute significantly to the measured *ex vivo* ERG bipolar cell component in P23H/*Gnat2^-/-^* mice and cannot explain the observed potentiation of *R_BC_* (Figure 4B-C, Table 1). Another possibility for the increased sensitivity of the rod bipolar cells in P23H mice is reduced inhibitory input into rod bipolar cells, *i.e.* disinhibition. To test this, we determined the contribution of GABA-, glycine-, and inhibitory dopaminergic signaling to the rod bipolar cell responses using *ex vivo* ERG in the retinas from *Gnat2^-/-^* and C57 wild-type mice (Figure 5B). No significant differences were found in the ERG a- or b-wave amplitudes between the control and treated conditions with any of the antagonists suggesting that disinhibition, as a mechanism, likely does not play a significant role in elevating rod bipolar cell responses observed in *ex vivo* ERG recordings from P23H mice (Figure 5B). Collectively, these results suggest that during rod degeneration and suppressed rod phototransduction triggered by the P23H rhodopsin mutation, the rod-rod bipolar cell signaling is strengthened via compensatory pathways that are most probably intrinsic to the rod axon terminal and/or rod bipolar cells.

**Figure 5.**
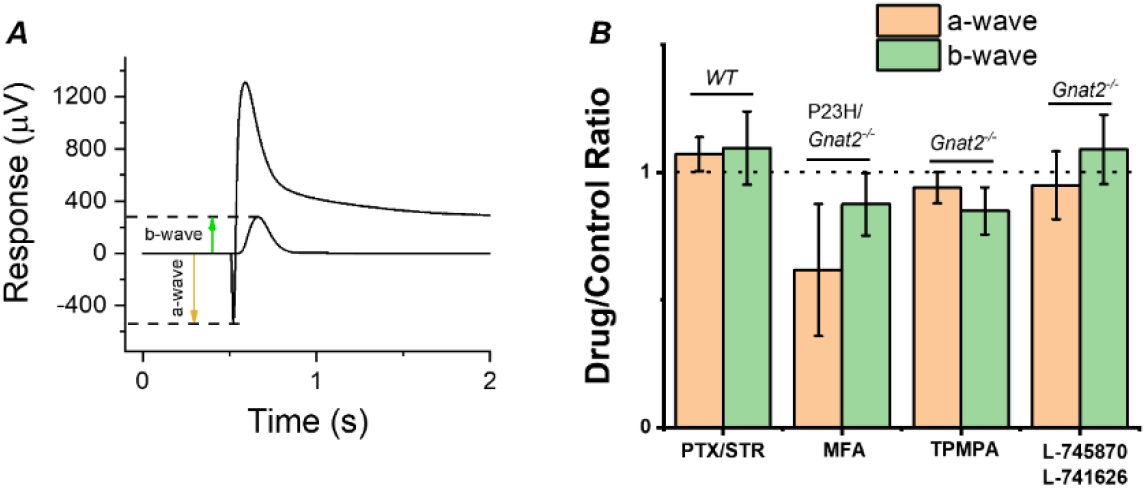
Contribution of gap junctions or inhibitory synaptic inputs to bipolar cells on the photoreceptor and bipolar cell components of the *ex vivo* ERG signal. (A) Example traces to dim and bright flashes recorded from a dark-adapted wild-type mouse retina perfused with Ames’ containing 100 μM BaCl_2_. B-wave amplitudes were measured from baseline using the dim flash responses and a-wave amplitudes from the baseline using the bright flash responses as indicated by arrows. (B) Mean α SEM a- (brown, a-wave_drug_/a-wave_control_) and b-wave (green, b-wave_drug_/b-wave_control_) ratios determined from the each individual retina perfused with control solution and solution containing 100 μM of the gap junction blocker meclofenamic acid (MFA), 100 μM of picrotoxin (PTX) and 10 μM strychnine (STR), 10 μM of the dopamine D2 and D4 receptor antagonists L-745870 and L-741626, respectively, or 50 μM the GABA_C_ receptor antagonist (1,2,5,6-Tetrahydropyridin-4-yl)methylphosphinic acid (TPMPA). Mouse genotypes for each drug experiment is indicated in Figure. No statistically significant changes in a- or b-wave amplitudes was observed with any of the drugs tested by paired t-test. n=6 retinas for PTX/STR, n=3 retinas for each MFA, TPMPA, and L-745870/L741626 experiments.

### Rod-mediated visual contrast sensitivity remains relatively normal despite loss of majority of rods

Our molecular genetics and functional analyses showed that compensatory pathways in the retina are expected to support retinal output even after 50-70% degeneration or functional decline of the rods in P23H mice. To test if these compensatory changes can promote vision, we applied an optomotor response test (OMR;(Prusky, Alam, Beekman, & Douglas, 2004) to evaluate behavioral visual contrast sensitivity (CS) in P23H/*Gnat2^-/-^* mice and their *Gnat2^-/-^* littermate controls (Figure 6A). This reflexive test was selected to avoid confounding factors caused by potential plasticity in the cortical brain areas that could also promote vision even when retinal output is suppressed (Koskela, Turunen, & Ala-Laurila, 2020). Our aim was to focus on the question of whether potentiation of the rod-rod bipolar cell signaling in P23H mice can help maintain high-sensitivity night vision. This goal was further facilitated by use of the *Gnat2^-/-^* background to abolish light responses of cones and the use of ambient light ranging from scotopic to photopic conditions. We found that CS decreased slightly in P23H/*Gnat2^-/-^* mice only at the dimmest background light up until 3-months of age in ambient light ranging from −3.3 to 0.85 log(Cd/m^2^) (Figure 6B,C). At 5-months of age, CS was suppressed at the two dimmest ambient light levels (−3.3 and −2.4 log(Cd/m^2^)), but not at brighter illuminances (Figure 6D). In contrast, a significant drop in performance occurred in 6-month-old P23H/*Gnat2^-/-^* mice at all except the brightest ambient light level (Figure 6D). A decrease in the sensitivity of rod-mediated vision was expected since absolute rod bipolar cell responses decreased in P23H mice as they aged beyond 1-month (Figures 3F and 4B) (Sarria et al., 2015). However, consistent with our finding that the rod bipolar cell sensitivity to their rod input was potentiated in P23H/*Gnat2^-/-^* mice up to 5-months of age, the decrease in scotopic contrast sensitivity was less severe than expected if based solely on the degeneration and functional decline of rod photoreceptors caused by the P23H rhodopsin mutation.

**Figure 6.**
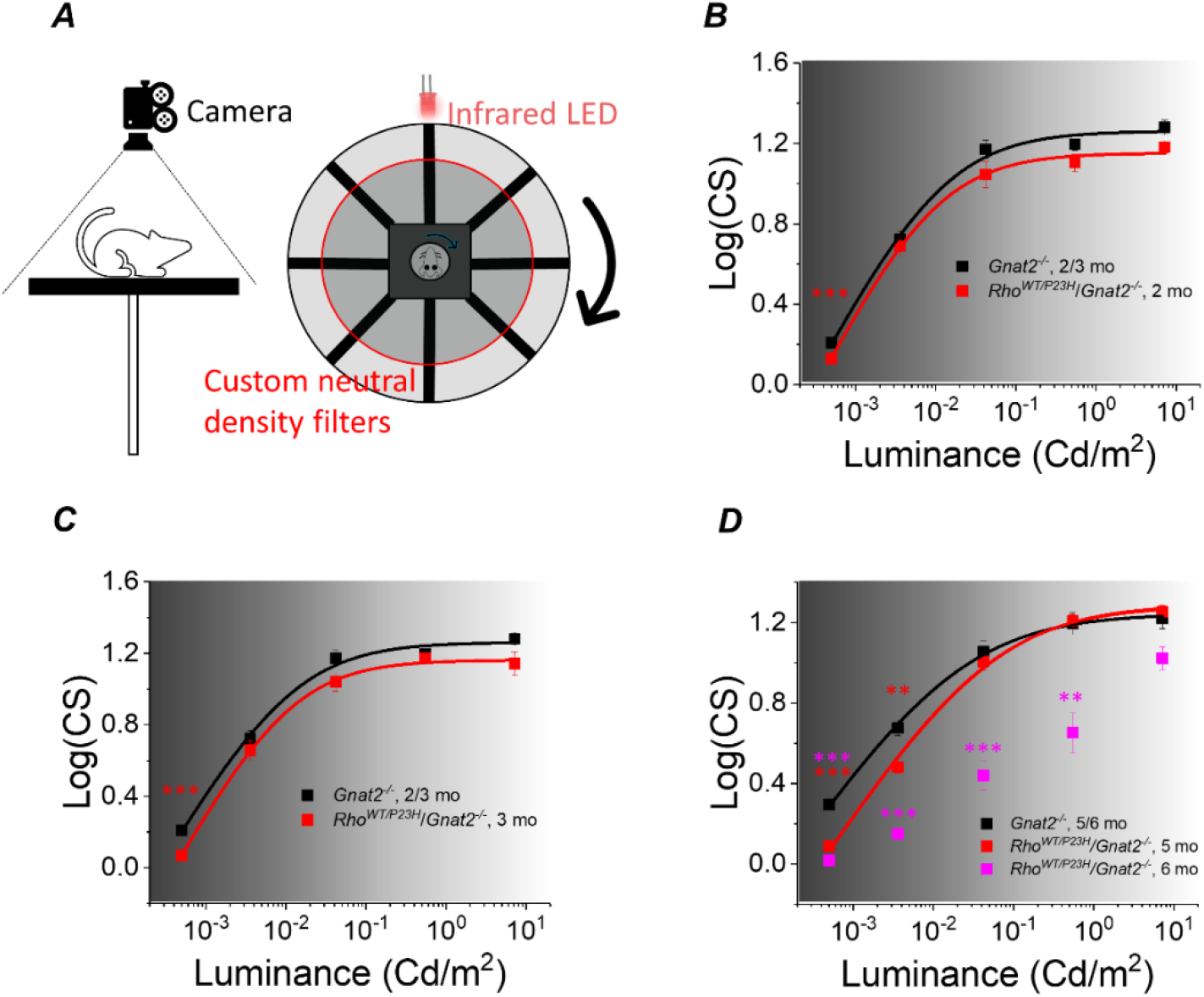
Visual contrast sensitivity (CS) mediated by rods is not severely compromised in P23H/*Gnat2^-/-^* mice up to five months of age. (A) Mouse contrast sensitivity threshold was tested using an optomotor reflex test and infrared visualization of the mouse head movement and cylindrical neutral density filters around the mouse platform. Contrast sensitivity was compared between *Rho^WT/P23H^*/*Gnat2^-/-^* and their age-matched *Gnat2^-/-^* littermate mice from dim scotopic up to mesopic/photopic conditions at 1 (B), 3 (C), and 5/6 (D) month of age. Smooth lines plot Hill-type Eq. (3) fitted to mean data points. \**P*<0.05, \*\**P*<0.005, \*\*\**P*<0.001, two-way ANOVA followed by Bonferroni’s posthoc test. (Control, 2-3 mo, n=9; P23H, 2 mo, n=11; P23H, 3 mo, n=6; control, 5-6 mo, n=9; P23H, 5 mo, n=4; P23H, 6 mo, n=5 mice)

## DISCUSSION

### P23H mice as a model to study the impact of early stage retinal remodeling on retinal function and vision

Retinal remodeling commonly refers to neuronal-glial changes in the retina in response to photoreceptor degeneration regardless of the etiology of the disease, and it is characterized by three distinct phases (Jones & Marc, 2005; Pfeiffer, Marc, & Jones, 2020). In phase 1, glial cells are activated, and neural reprogramming begins due to photoreceptor death, typically starting in rods that mediate our night vision. Phase 2 refers to an advanced disease stage when collateral, secondary cone cell death ensues. Phase 3 is a terminal phase and is thought to begin when all the photoreceptors have degenerated. According to this classification, our study using an adRP mouse model begins when remodeling is in phase 1 and ends in the earliest steps of phase 2 (Figure 1).

We employed knock-in mice that carry a heterozygous P23H rhodopsin mutation. The P23H mouse line is a well characterized model for the most common form of adRP, and closely parallels the pathological characteristics of the human disease (Sakami et al., 2014; Sakami et al., 2011). Typically, the clinical disease first manifests as night blindness followed by a gradual loss of visual field (Berson, Rosner, Sandberg, & Dryja, 1991). However, the disease severity and progression are moderate compared to many other types of RP, and the patients can retain normal visual acuity well into their 30s (Berson et al., 1991). This may be due to the cone population preserved in many patients (Hartong, Berson, & Dryja, 2006). We found that the number of cone cells remained normal in P23H mice up to 5-months of age, the last time point studied (Figure 1K). Cone function was normal in 1- and 3-month-old P23H mice but significantly decreased at 5-months of age when compared to their WT littermates (Figure 1L). An earlier study demonstrated that the maximal rod response amplitude is suppressed by only 5% at post-natal day (P) 14 in heterozygous P23H mice (Sakami et al., 2014), implying practically normal amplitudes at eye opening. We found that the maximal ERG response arising from the rod population was reduced by 45 – 60% in 1-month-old P23H mice compared to control mice depending on the method of recording (*in vivo* vs. *ex vivo*) and the background of the mice (C57 vs. *Gnat2^-/-^*) (Figures 1J, 3D and 4A). However, the waveform of the ERG response appeared qualitatively normal, and the first positive deflection of the *in vivo* ERG signal (b-wave) (Figure 1I) as well as the rod bipolar cell responses derived using *ex vivo* ERG (Figure 3F and 4B, Table 1) were comparable between 1-month-old P23H and control mice. Therefore, we concluded that 1-month of age represents an optimal time for a molecular genetics screen of P23H mice as by this age a functional deficit of the rod population is significant, whereas their degeneration is not yet severe (Figures 1, 3 and 4). Furthermore, at 1-month of age the confounding factors arising from any developmental plasticity associated with the retina is avoided due to the full maturation of the mouse retina at this age (Young, 1985).

We first hypothesized that if homeostatic plasticity occurs in response to rod cell death in the retina, it must show a discernible fingerprint in the retinal transcriptome. As expected, due to a loss of rods, several photoreceptor selective gene clusters were downregulated in P23H retinas (Figure 2B). In contrast, pronounced upregulation arose in pathways involving neuronal development, such as synaptic organization, axonogenesis and cell adhesion (Figure 2C, D). We also found several enriched pathways that overlap between cell stress responses and neuronal plasticity, such as the MAPK, NF-kappa B and TNF signaling pathways (Pozniak, White, & Khalili, 2014). In the case of P23H retinas, it is difficult to determine whether these major transcription regulators are upregulated primarily due to cell degeneration and stress or due to induction of plasticity machinery. Nevertheless, our RNA-seq analysis is consistent with our hypothesis that a robust neuronal network adaptation in the retina takes place in the early stages of photoreceptor degenerative disease caused by the P23H rhodopsin mutation. At the same age, we found that while the photoreceptor component of the *ex vivo* ERG response was significantly suppressed (Figures 3E and 4A), the light sensitivity of the rod bipolar cell responses was similar to that in control mice (Figures 3F and 4B, Table 1). In the C57 background, the light-sensitivity of bipolar cells tended to be even higher in 1-month-old P23H mice as compared to WT controls (Figure 3F, Table 1). However, this may be due to the contribution from increased light responses of cones observed *in vivo* (Figure 1L), since without the contribution of cone phototransduction, the rod bipolar cell sensitivity slightly decreased in P23H/*Gnat2^-/-^* mice (Figure 4B, Table 1). However, the compensatory sensitization of the rod bipolar cells to their rod input cannot be explained simply by upregulation of cone-mediated signaling, since the same phenomenon was observed in P23H mice both in C57 and *Gnat2^-/-^* backgrounds (Figures 4C, Table 1). The P23H/*Gnat2^-/-^* mice were also observed to perform almost normally until 3-months of age in a behavioral visual contrast threshold task, even at the lowest luminance (Fig. 5). At this age, the overall rod input already decreased by more than half in P23H mice which was expected to significantly elevate contrast thresholds (or decrease contrast sensitivity) specifically at the two lowest luminance levels where the contrast sensitivity is proportional to background light intensity (Figures 3E and 4A). Therefore, it appears that the gradual loss of the rod photoreceptors, or declined input from them, triggers functional homeostatic adaptation in the retina that allows maintenance of visual function downstream of the photoreceptors.

### Is retinal remodeling and plasticity supporting or impairing retinal output and vision?

Our results using the P23H mouse model appear at odds with some prior research casting a rather negative connotation over retinal remodeling following the loss of photoreceptors. Such studies proposed that retinal remodeling exacerbates vision loss and therefore may interfere with strategies to restore vision in the blind (Jones et al., 2003; Marc, Jones, Watt, & Strettoi, 2003; Telias et al., 2019; Toychiev et al., 2013). What might underlie this apparent discrepancy with our findings? We speculate that the contrasting conclusions could stem from the retinal remodeling phase at which various retinal degeneration models with significantly different rates of photoreceptor degeneration have been studied. The list of retinal degeneration mouse models in which remodeling has been investigated is extensive; however, currently the most utilized models according to the literature are *rd1* and *rd10* mice in which photoreceptors degenerate rapidly due to mutations of phosphodiesterase 6 (*Pde6*) in rods. The *rd1* mouse model represents an extremely severe disease where rod death starts at P8 and progresses to practically complete loss of all photoreceptors by 4 weeks of age. The *rd10* model exhibits somewhat slower progression of the disease. Their dark-adapted ERG response is observable but already significantly compromised at P18 and by 1-month of age the light responses of rods are practically abolished and most of their photoreceptors have degenerated, particularly in the central retina (Chang, Hawes, Pardue, et al., 2007; Gargini, Terzibasi, Mazzoni, & Strettoi, 2007; Wang et al., 2018). Thus, it is possible that functional and behavioral studies using *rd1* and *rd10* mice, typically conducted with mice older than 1-month of age, may have missed the window for observing constructive remodeling expected to promote retinal output and vision. Instead, the heterozygous P23H animal model provides a useful window of opportunity for investigating the impact of photoreceptor degeneration on early stage retinal remodeling (phase 1).

Recent studies using the specific deletion of ~50 – 90% of rod bipolar cells or cones during development or in mature mice have revealed homeostatic mechanisms associated with the maintenance of normal retinal output and vision (Care et al., 2019; Johnson et al., 2017; Shen et al., 2020; Tien, Soto, & Kerschensteiner, 2017). Here we used a P23H knock-in mouse model of RP that recapitulates the human disease. Our results are consistent with the idea that at least during phase 1 of remodeling, photoreceptor death induces compensatory pathways that support close to normal retinal output and rod-mediated vision (Figs. 2, 4, and 6). Even at the stage where we observed prominent cone dysfunction (Fig. 1L) in 5- or even 6-month-old P23H/*Gnat2^-/-^* mice, the rod bipolar cells continued to increase their sensitivity to rod input (Figure 4C-F, Table 1), although at this point the light responses of rods were already small (Figure 4A) and the absolute amplitudes of the rod bipolar cell light responses had significantly decreased (Figure 4B) together with lower contrast sensitivity of their rod-mediated vision in dim light (Figure 6D). These results therefore suggest that up to a point where some residual rod light responses exist, the retina can adapt towards supporting normal retinal output and scotopic vision. In this manner, patients at this stage of retinal degenerative disease might be expected to benefit from various treatments designed to restore photoreceptors and their light responses. However, the duration during which the inner retina remains capable of processing light signals and therefore is amenable to these types of treatments remains unknown. Studies with other animal models suggest that corruption of the inner retina would become a problem at later time points, including in P23H mutation-associated RP models. Quantitative functional and behavioral studies at later stages will be needed to address this question.

### Mechanism for increased sensitivity of rod bipolar cells to their rod input

Our RNA-seq and electrophysiology data strongly suggest that homeostatic plasticity is induced at early stages of rod degeneration in the P23H mouse retina. Still, the homeostatic plasticity mechanisms for the increased sensitivity of rod bipolar cells to their rod input remain unclear. Possibilities include 1) formation of new synapses between rod bipolar cells and surviving photoreceptors, 2) increased electrical coupling of rods to the cone pathway, 3) reduced inhibitory drive to rod bipolar cells (disinhibition), 4) pre- and/or postsynaptic scaling, and 5) increased excitability through axonal remodeling of rod bipolar cells. We tested hypotheses 2 and 3 by using blockers for gap junctions and major inhibitory postsynaptic receptors (Figure 5). Our findings did not support either of these hypotheses, and it appears that the potentiation of the ERG b-wave is not mediated by disinhibition or altered gap junctional coupling in P23H mice. The first hypothesis, on the other hand, aligns with recent studies showing that cone bipolar cell dendrites formed new synapses with surviving cones after ~50% of cones had been ablated in juvenile or mature mice (Care et al., 2019; Shen et al., 2020). Interestingly, ablation of 50 – 90% of rod bipolar cells during development led to an extension of the dendrites of the remaining rod bipolar cells so that they contacted more rods (Johnson et al., 2017). These homeostatic mechanisms appeared to support normal retinal function and vision. It also has been shown that restoration of rod function in juvenile mice, which lacked rod function during development, can lead to the formation of new functional synapses between rods and rod bipolar cells (Wang et al., 2019). Thus, it is possible that photoreceptor degeneration leads to the formation of new synapses between rod bipolar cells and surviving rods and/or an increase in the number of postsynaptic densities adjacent to individual rod terminals (see (Johnson et al., 2017). Our RNA-seq data show upregulation in a number of ion channels and G protein-coupled receptors, such as the *Trpm1* cation channel crucial for ON bipolar cell depolarization as well as glutamatergic receptors that could contribute either to the strengthening of individual rod-rod bipolar cell synapses or increased excitability of the rod bipolar cells. Additional work will be needed to resolve the exact homeostatic plasticity mechanisms that promote rod-rod bipolar cell signaling and visual function. Such knowledge could then help us understand how sensory systems respond to potentially compromising circumstances throughout life, and as well help in designing interventions to enhance homeostatic plasticity when needed.

## MATERIAL & METHODS

### Animals

Initially, we crossed *Rho^P23H/P23H^* males (Sakami et al., 2011) with C57Bl/6J mice (The Jackson Laboratory, stock # 000664) to generate mice carrying heterozygous P23H mutation in rhodopsin. Experimental (*Rho^P23H/WT^*) and littermate control mice were generated using *Rho^P23H/WT^* and *Rho^WT/WT^* breeding pairs. We also generated a P23H mouse line on *Gnat2^-/-^* (Ronning et al., 2018) background by crossing P23H mice with *Gnat2^-/-^* mice, a kind gift from Dr. Marie Burns, University of California, Davis. Experimental (*RhoP^23H/WT^*/*Gnat2^-/-^*) and littermate controls (*Gnat2^-/-^*) were generated using *Rho^P23H/WT^*/*Gnat2^-/-^* and *Rho^WT/WT^*/*Gnat2^-/-^* breeding pairs. Both male and female mice were used in this study. Mice were kept under 12/12 h light cycle with free access to food and water. All experimental protocols adhered to Guide for the Care and Use of Laboratory Animals and were approved by the institutional Animal Studies Committees at the University of Utah and University of California, Irvine.

### Histology

After euthanasia, the superior side of each mouse eye was marked with a thread burner and then enucleated. Eyes were then fixed in Hartman’s fixative (Sigma Aldrich, St. Louis, MO) for at least 24 hours. The eyes were kept in 70 % ethanol before embedding into paraffin blocks. Sections were cut at 10 μm thickness in a nasal-temporal orientation. Retinal panorama images were captured using a light microscope. Representative magnified images shown in Figure 1B,C were taken at the superior middle retina centered at ~700 μm from the optic nerve head (ONH).

### Optical coherence tomography

Mice were anesthetized with ketamine (100 mg/kg, KetaVed®, Bioniche Teoranta, Inverin Co, Galway, Ireland) and xylazine (10 mg/mg, Rompun®, Bayer, Shawnee Mission, KS) by intraperitoneal injection, and their pupils were dilated with 1% tropicamide. OCT was performed with a Bioptigen spectral-domain OCT device (Leica Microsystems Inc., Buffalo Grove, IL). Four frames of OCT b-scan images were acquired from a series of 1200 a-scans. ONL thickness was measured 500 μm from the ONH at nasal, temporal, superior and inferior quadrants. The ONL thickness was averaged over the four retinal quadrants, and this average was used in the analysis.

### Preparation of retinal whole mounts and cone counting

Retina whole mounts for cone population analyses were performed as described earlier (Leinonen, Choi, Gardella, Kefalov, & Palczewski, 2019). In brief, the superior side of each eye was marked with a permanent marker before enucleation. Eyes were then fixed in 4 % paraformaldehyde in isotonic phosphate buffered saline (PBS) for 1 hour after enucleation, and the retina was dissected away and processed as a whole-mount sample and subsequently stained using polyclonal goat S-opsin (1:1000 dilution, Bethyl Laboratories, Montgomery, TX) and polyclonal rabbit M-opsin (1:1000 dilution, Novus Biologicals, Littleton, CO) primary antibodies and fluorescent secondary antibodies (dilution for both 1:500; donkey anti-goat Alexa Fluor 488 and donkey anti-rabbit Alexa Fluor 647; Abcam, Cambridge, UK). The whole retinal area was imaged using a fluorescence light-microscope (Leica DMI6000B, Leica Microsystems Inc, Buffalo Grove, IL) equipped with an automated stage, a 20× objective, and green (excitation 480/40 nm) and far red (excitation 620/60 nm) fluorescence filter channels. Cone counts were computed from each individual image using MetaMorph 7.8 software (Molecular Devices, Sunnyvale, CA) and summed together to yield a whole retina count. For illustrative panorama images, the individual images were stitched together using MetaMorph 7.8 software.

### *In vivo* electroretinography (ERG)

*In vivo* ERG was performed as previously described (Orban et al., 2018) using a Diagnosys Celeris rodent ERG device (Diagnosys, Lowell, MA). Briefly, a mouse was anesthetized with ketamine & xylazine and placed on a heating pad at 37 °C, and its pupils were moistened with 2.5% hypromellose eye lubricant (HUB Pharmaceuticals, Rancho Cucamonga, CA). Light stimulation was produced by an in-house scripted stimulation series in Espion software (version 6; Diagnosys). Before scotopic recordings, the mice were dark-adapted overnight and animal handling before recording was performed under dim red light. The eyes were stimulated with a green LED (peak 544 nm, bandwidth 160 nm) using a 13-step ascending flash intensity series ranging from 0.00015 to 300 cd·s/m^2^.

Before photopic ERG recordings, mice were kept in a vivarium. After induction of anesthesia, the eyes were first adapted to a rod-suppressing green background light at 20 cd/m^2^ for 1 min. Stimulation was performed with a blue LED (peak emission 460 nm, bandwidth ∼100 nm) at light intensity increments of 0.1, 1.0, and 10.0 cd·s/m^2^. As the short wavelength cone opsin (S-opsin) and medium wavelength cone opsin (M-opsin) sensitivities peak at 360 and 508 nm in mice (Nikonov, Kholodenko, Lem, & Pugh, 2006), respectively, the blue LED stimulates both S- and M-opsins. LED light emission spectra were measured with a Specbos 1211UV spectroradiometer (JETI Technische Instrumente GmbH, Jena, Germany).

The ERG signal was acquired at 2 kHz and filtered with a low frequency cutoff at 0.25 Hz and a high frequency cutoff at 300 Hz. Espion software automatically detected the ERG a-wave (first negative ERG component) and b-wave (first positive ERG component).

### RNA isolation and RNA-sequencing

Mice were euthanized by cervical dislocation, and eyes were promptly harvested. Retinas were dissected away from the eyecup and put into RNAlater solution (Qiagen, Germantown, MD). The samples were stored at −80 °C before processing. The retinas were homogenized using a motorized pestle, and the homogenate was spun in a tabletop centrifuge at full speed for 3 min. Supernatant was collected and RNA extracted using a Qiagen RNeasy Mini Kit following the manufacturer’s instructions. On-column DNAase digestion (Qiagen RNase-Free DNase) was used to remove any genomic DNA contamination from the sample. Five hundred ng of purified RNA were used for cDNA synthesis using an iScript cDNA synthesis kit (Bio-Rad, Irvine, CA), and real-time quantitative PCR (qPCR) was used as one of the initial sample quality checks before RNA-sequencing (RNA-seq).

Final RNA quality control (QC), library preparation, and RNA-seq were performed by Sacramento NovoGene Co., Ltd. (Sacramento, CA) using the Illumina HiSeq instrument. QC was performed with agarose gel electrophoresis and Nanodrop and Agilent 2100 devices. The gene expression data were aligned with a reference genome (mus musculus mm10) using Tophat2 (http://ccb.jhu.edu/software/tophat/index.shtml). The gene expression level was measured by transcript abundance and reported as Fragments Per Kilobase of transcript sequence per Millions base pairs (FPKM) sequenced. As the first measure of dataset reliability, a Pearson correlation was conducted between all samples. Subsequently, to test data integrity between experimental groups (WT female, n=4; WT male, n=3; P23H female, n=4; P23H male, n=3), the FPKM distribution was inspected in violin plots. Next, Venn diagrams were inspected to reveal relationships between the four groups, after which the sexes were pooled and only genotypes (WT females+males vs P23H females+males) were compared in further analyses. The Gene Ontology (GO) and KEGG analyses, and predicted protein-protein interaction analysis using the STRING database (https://string-db.org/cgi/input.pl), were obtained to reveal gene expression changes at the level of networks. Gene expression differences of single genes between genotypes was reported as log2(mean of P23H/mean of WT) and an adjusted *P*-value of <0.05 was used as a level of statistical significance.

### Immunoblotting

The mice were euthanized by cervical dislocation, eyes were promptly harvested, and retinas were quickly dissected away from the eyecup and immediately frozen using liquid nitrogen. A single retina extract was homogenized in ice-cold lysis buffer (70 μl) containing 100 mM Tris-HCl, 10 mM magnesium acetate, 6 M urea, 2 % SDS, benzonase nuclease (25 U per ml), protease inhibitor (cOmplete Mini EDTA-free, Roche Diagnostics, Indianapolis, IN) and phosphate inhibitor cocktails I and II (Sigma Aldrich). Homogenates were centrifuged and the supernatants mixed with Laemlli sample buffer (Bio-Rad). Protein concentration in each sample was determined using a Pierce BCA protein assay kit (ThermoFisher, Waltham, MA). Forty-eight μg of protein were loaded onto a Mini-Protean TGX gel (Bio-Rad) and electrophoresis was run on an ice-bed. Proteins were transferred to a nitrocellulose membrane. The membrane was blocked with 5 % fat-free milk in Tris-buffered saline containing 0.1 % Tween 20 (TBST) for 1 h, and thereafter incubated in primary antibody in 1 % milk in TBST. After an overnight incubation, the membrane was washed three times with TBST and thereafter incubated in fluorescent secondary antibody solution (LI-COR, Lincoln, NE) at 1:10 000 dilution for 2 h. The membrane was washed three times with TBST and subsequently imaged using an Odyssey imager (LI-COR). The primary antibodies used were mouse anti-phospho-ERK1/2 (dilution 1:1000), rabbit anti-ERK1/2 (dilution 1:2000) and rabbit α-tubulin (dilution 1:2000) all purchased from Cell Signaling Technology (Danvers, MA). The phospho-ERK1/2 expression was evaluated first using a 680-nm secondary anti-mouse antibody, and the same membrane was re-incubated later in ERK1/2 and α-tubulin solution using an 800-nm secondary anti-rabbit antibody.

### E*x vivo* Electroretinogram

Transretinal (*ex vivo*) electroretinography (ERG) was conducted as described previously (Vinberg et al., 2014; Vinberg, Wang, Molday, Chen, & Kefalov, 2015). Briefly, mice were dark adapted for at least two hours and euthanized by CO_2_ and cervical dislocation under dim red light. Eyes were enucleated and immediately placed into carbonated (95% O_2_/5% CO_2_) Ames’ medium (A1420; Sigma-Aldrich, St. Louis, MO) containing 1.932 g/L NaHCO_3_ (Sigma-Aldrich, St. Louis, MO). The retina was dissected from the eye and transferred to the recording chamber which was perfused at a rate of 1 mL/min with heated (37 °C) Ames’ medium containing 100 μM BaCl2 (Sigma-Aldrich, St. Louis, MO) to remove the Müller glia contribution to the ERG signal (Bolnick et al., 1979). To isolate the photoreceptor component of the ERG signal, 40 μM DL-AP4 (Cat. #0101; Tocris Bioscience) was added to the perfusion medium. In some experiments (Figure 5), the following drugs were added into Ames’ (μM; supplier Sigma-Aldrich, St. Louis, MO unless otherwise noted): 100 picrotoxin (PTX) to block GABA receptors, 10 strychnine (STR) to block glycine receptors, 100 meclofenamic acid (MFA) to block electrical coupling via gap junctions, 50 (1,2,5,6-Tetrahydropyridin-4-yl)methylphosphinic acid (TPMPA) to block GABAC receptors, and 10 L-741626 and L-745870 (both from Tocris Bioscience) to block D_2_ and D_4_ receptors respectively. Drugs were first dissolved in ethanol (PTX), DMSO (L-741626) or water (STR, L-745870, MFA, TPMPA), and 0.02% −0.2% of the stock solutions were added into Ames’ to achieve the final concentration indicated above. For control solution, the same amount of water, DMSO or ethanol instead of the drug-containing stock solution was added.

From the dark-adapted retina, ERG responses to flashes of light (505 nm, 2-10 ms flash duration) from ~17 photons μm^-2^ up to ~20,000 photons μm^-2^ were recorded (10 trials/dimmer flash intensity and 5 trials/brighter flashes). ON bipolar cell responses were determined by subtracting the photoreceptor response (containing DL-AP4) from the ERG response without DL-AP4. Photoreceptor response and ON bipolar cell response amplitudes were measured and their ratio was calculated and plotted as a function of light flash intensity (see Figure 3). To determine flash intensity (*I*) required to generate half-maximal photoreceptor or ON bipolar cell responses (*I_1/2_*), a Hill function

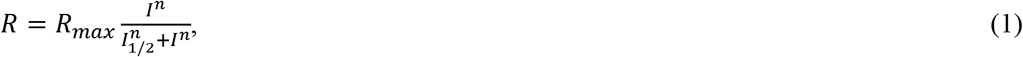

was fitted to amplitude (*R*) data using the Originlab’s (2018, build: 9.5.0193) nonlinear curve fit tool. *R_max_* is the maximal response amplitude to a bright flash (set based on data) and n is a steepness factor that was set to 1 for photoreceptor responses but was let to vary freely to produce optimal fits to *R_BC_* (bipolar cell response amplitude) data. We also fit similar Hill functions to data plotting *R_BC_* as a function of *R_PR_*:

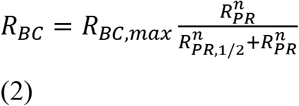

to determine the photoreceptor input required to generate half-maximal ON bipolar cell response, *R_PR,1/2_*.

### Measurement of visual contrast sensitivity

An optomotor reflex test (OMR) was used to determine contrast sensitivity threshold using a commercially available OptoMotry^©^ system (CerebralMechanics Inc., Canada) (Prusky et al., 2004) customized with an infrared LED in the chamber for visualization of mouse head movement by a camera attached to the lid of the box (Figure 6A). Additional cylindrical neutral density filters (in layers) were placed around the mouse platform to assess contrast sensitivity in scotopic and mesopic-photopic conditions. The ambient luminance (Cd/m^2^) was calibrated using an optometer (UDTi FlexOptometer, photometric sensor model 2151, Gamma Scientific, CA, USA). Since the dynamic range of the sensor did not reach the dimmest conditions, those were derived based on separate measurements of the attenuation factor for each neutral density filter layer. Before each experiment mice were dark-adapted overnight, and the testing started at the dimmest light condition. The test was administered in a single-blinded fashion, with the experimenter unaware of the stimulus presented to the mouse. Horizontally drifting vertical sine-wave gratings moving left or right with different contrast were presented at a spatial frequency of 0.128 cycles/degree and a grating rotation speed (drift speed) of 5.4 degrees/s to measure contrast thresholds (CT, 1 – 100%) at different ambient luminance (*L*) levels from 5*10^-4^ to 7 Cd/m^2^. Contrast sensitivity (CS) is defined as 1/CT and was plotted as a function of ambient luminance. A Hill function defined by

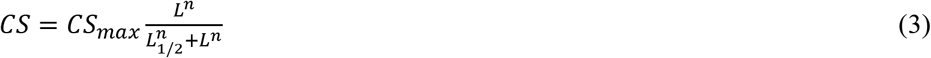

was fit to the mean luminance-CS data with *L_1/2_* (luminance at which *CS* = 0.5*CS*_max_), *CS*_max_ (maximal CS at ~1-7 Cd/m^2^) and n (steepness factor) as free parameters.

### Statistical analysis

ONL thickness analysis was tested using one-way ANOVA. Repeated measures two-way ANOVA was performed for *in vivo* ERG stimuli series comparisons. Two-way ANOVA was used to analyze OMR and R_BC_/R_PR_ data at various luminance/intensity levels. The ANOVAs were followed by Bonferroni’s post hoc tests. *Ex vivo* ERG parameters from fitted functions or measured *Rmax* between control and P23H mice were analyzed by a two-tailed t-test. A paired two-tailed t-test was used to determine if inhibitors affected the ERG a- or b-waves (Figure 5). Results are displayed as mean± SEM and statistical significance was set at *P*<0.05.

## Supporting information

Fig. 2. - Supplement 1. Activation of mitogen-activated protein kinase (MAPK) pathway in 1-mo P23H retinas. T-test: **P<0.01, ***P<0.001.

## Acknowledgments

H.L. was supported by Fight for Sight, Eye and Tissue Bank Foundation (Finland), the Finnish Cultural Foundation and the Orion Research Foundation. F.V. was supported by NIH grant EY026651, the International Retinal Research Foundation, and Research to Prevent Blindness / Dr. H. James and Carole Free Career Development Awards. K.P was supported by NIH grants EY009339 and R24 EY027283. K.P is the Irving H. Leopold Chair of Ophthalmology. The authors also acknowledge support from an RPB unrestricted grant to the Department of Ophthalmology, University of California, Irvine and to the Department of Ophthalmology and Visual Sciences, University of Utah. We thank Dr. Marie Burns (University of California, Davis) for providing the *Gnat2^-/-^* mouse strain used in this study.

## Author contributions

H.L.: Conceptualization, Data curation, Formal analysis, Funding acquisition, Supervision, Validation, Investigation, Visualization, Methodology, Writing - original draft, Writing - review and editing.

N.C.P.: Investigation, Visualization, Formal analysis, Writing - review and editing.

T.B.: Investigation.

J. S.: Investigation.

K. P.: Resources, Funding acquisition, Supervision, Writing - review and editing.

F.V.: Conceptualization, Resources, Data curation, Formal analysis, Supervision, Funding acquisition, Validation, Investigation, Visualization, Methodology, Project administration, Writing - original draft, Writing - review and editing.

All authors read, reviewed, and accepted the final manuscript.

## Abbreviations

adRP: autosomal dominant Retinitis Pigmentosa
AMD: Age-Related Macular Degeneration
ANOVA: analysis of variance
CS: contrast sensitivity
ERG: electroretinography
GO: gene ontology
IS: inner segment
KEGG: Kyoto Encyclopedia of Genes and Genomes
MAPK: mitogen-activated protein kinase
DMSO: Dimethyl sulfoxide
PTX: picrotoxin
STR: strychnine
MFA: meclofenamic acid
NF-kappa B: nuclear factor kappa B
OCT: optical coherence tomography
OMR: optomotor response
ONH: optic nerve head
ONL: outer nuclear layer
OS: outer segment
PBS: phosphate-buffered saline
RP: Retinitis Pigmentosa
TBS: Tris-buffered saline
TNF: tumor necrosis factor
WT: wild-type.

## Notes

**Conflict of interests:** None

### Competing Interest Statement

The authors have declared no competing interest.

## REFERENCES

Abraham, W. C., & Bear, M. F. (1996). Metaplasticity: the plasticity of synaptic plasticity. Trends Neurosci, 19(4), 126–130. doi:10.1016/s0166-2236(96)80018-x

Aleman, T. S., LaVail, M. M., Montemayor, R., Ying, G., Maguire, M. M., Laties, A. M., … Cideciyan, A. V. (2001). Augmented rod bipolar cell function in partial receptor loss: an ERG study in P23H rhodopsin transgenic and aging normal rats. Vision Res, 41(21), 2779–2797.

Beier, C., Hovhannisyan, A., Weiser, S., Kung, J., Lee, S., Lee, D. Y., … Sher, A. (2017). Deafferented Adult Rod Bipolar Cells Create New Synapses with Photoreceptors to Restore Vision. J Neurosci, 37(17), 4635–4644. doi:10.1523/JNEUROSCI.2570-16.2017

Beier, C., Palanker, D., & Sher, A. (2018). Stereotyped Synaptic Connectivity Is Restored during Circuit Repair in the Adult Mammalian Retina. Curr Biol, 28(11), 1818–1824 e1812. doi:10.1016/j.cub.2018.04.063

Beltran, W. A. (2009). The use of canine models of inherited retinal degeneration to test novel therapeutic approaches. Vet Ophthalmol, 12(3), 192–204. doi:10.1111/j.1463-5224.2009.00694.x

Berson, E. L., Rosner, B., Sandberg, M. A., & Dryja, T. P. (1991). Ocular findings in patients with autosomal dominant retinitis pigmentosa and a rhodopsin gene defect (Pro-23-His). Arch Ophthalmol, 109(1), 92–101. doi:10.1001/archopht.1991.01080010094039

Bolnick, D. A., Walter, A. E., & Sillman, A. J. (1979). Barium suppresses slow PIII in perfused bullfrog retina. Vision Res, 19(10), 1117–1119. doi:10.1016/0042-6989(79)90006-3

Burrone, J., & Murthy, V. N. (2003). Synaptic gain control and homeostasis. Curr Opin Neurobiol, 13(5), 560–567. doi:10.1016/j.conb.2003.09.007

Care, R. A., Kastner, D. B., De la Huerta, I., Pan, S., Khoche, A., Della Santina, L., … Dunn, F. A. (2019). Partial Cone Loss Triggers Synapse-Specific Remodeling and Spatial Receptive Field Rearrangements in a Mature Retinal Circuit. Cell Rep, 27(7), 2171–2183 e2175. doi:10.1016/j.celrep.2019.04.065

Chang, B., Hawes, N. L., Davisson, M. T., & Heckenlively, J.R.. (2007). Mouse Models of RP. In J. Tombran-Tink & C.J. Barnstable (Eds.), Retinal Degenerations: Biology, Diagnostics, and Therapeutics. Totowa, NJ: Humana Press.

Chang, B., Hawes, N. L., Pardue, M. T., German, A. M., Hurd, R. E., Davisson, M. T., … Boatright, J. H. (2007). Two mouse retinal degenerations caused by missense mutations in the beta-subunit of rod cGMP phosphodiesterase gene. Vision Res, 47(5), 624–633. doi:10.1016/j.visres.2006.11.020

Desai, N. S., Cudmore, R. H., Nelson, S. B., & Turrigiano, G. G. (2002). Critical periods for experience-dependent synaptic scaling in visual cortex. Nat Neurosci, 5(8), 783–789. doi:10.1038/nn878

Fariss, R. N., Li, Z. Y., & Milam, A. H. (2000). Abnormalities in rod photoreceptors, amacrine cells, and horizontal cells in human retinas with retinitis pigmentosa. Am J Ophthalmol, 129(2), 215–223.

Gainey, M. A., & Feldman, D. E. (2017). Multiple shared mechanisms for homeostatic plasticity in rodent somatosensory and visual cortex. Philos Trans R Soc Lond B Biol Sci, 372(1715). doi:10.1098/rstb.2016.0157

Gargini, C., Terzibasi, E., Mazzoni, F., & Strettoi, E. (2007). Retinal organization in the retinal degeneration 10 (rd10) mutant mouse: a morphological and ERG study. J Comp Neurol, 500(2), 222–238. doi:10.1002/cne.21144

Goel, A., Jiang, B., Xu, L. W., Song, L., Kirkwood, A., & Lee, H. K. (2006). Cross-modal regulation of synaptic AMPA receptors in primary sensory cortices by visual experience. Nat Neurosci, 9(8), 1001–1003. doi:10.1038/nn1725

Goel, A., & Lee, H. K. (2007). Persistence of experience-induced homeostatic synaptic plasticity through adulthood in superficial layers of mouse visual cortex. J Neurosci, 27(25), 6692–6700. doi:10.1523/JNEUROSCI.5038-06.2007

Goo, Y. S., Ahn, K. N., Song, Y. J., Ahn, S. H., Han, S. K., Ryu, S. B., & Kim, K. H. (2011). Spontaneous Oscillatory Rhythm in Retinal Activities of Two Retinal Degeneration (rd1 and rd10) Mice. Korean J Physiol Pharmacol, 15(6), 415–422. doi:10.4196/kjpp.2011.15.6.415

Hartong, D. T., Berson, E. L., & Dryja, T. P. (2006). Retinitis pigmentosa. Lancet, 368(9549), 1795–1809. doi:10.1016/S0140-6736(06)69740-7

Haverkamp, S., Michalakis, S., Claes, E., Seeliger, M. W., Humphries, P., Biel, M., & Feigenspan, A. (2006). Synaptic plasticity in CNGA3(-/-) mice: cone bipolar cells react on the missing cone input and form ectopic synapses with rods. J Neurosci, 26(19), 5248–5255. doi:10.1523/JNEUROSCI.4483-05.2006

Johnson, R. E., Tien, N. W., Shen, N., Pearson, J. T., Soto, F., & Kerschensteiner, D. (2017). Homeostatic plasticity shapes the visual system’s first synapse. Nat Commun, 8(1), 1220. doi:10.1038/s41467-017-01332-7

Jones, B. W., & Marc, R. E. (2005). Retinal remodeling during retinal degeneration. Exp Eye Res, 81(2), 123–137. doi:10.1016/j.exer.2005.03.006

Jones, B. W., Pfeiffer, R. L., Ferrell, W. D., Watt, C. B., Marmor, M., & Marc, R. E. (2016). Retinal remodeling in human retinitis pigmentosa. Exp Eye Res, 150, 149–165. doi:10.1016/j.exer.2016.03.018

Jones, B. W., Pfeiffer, R. L., Ferrell, W. D., Watt, C. B., Tucker, J., & Marc, R. E. (2016). Retinal Remodeling and Metabolic Alterations in Human AMD. Front Cell Neurosci, 10, 103. doi:10.3389/fncel.2016.00103

Jones, B. W., Watt, C. B., Frederick, J. M., Baehr, W., Chen, C. K., Levine, E. M., … Marc, R. E. (2003). Retinal remodeling triggered by photoreceptor degenerations. J Comp Neurol, 464(1), 1–16. doi:10.1002/cne.10703

Koskela, S., Turunen, T., & Ala-Laurila, P. (2020). Mice Reach Higher Visual Sensitivity at Night by Using a More Efficient Behavioral Strategy. Curr Biol, 30(1), 42–53 e44. doi:10.1016/j.cub.2019.11.021

LaVail, M. M., Nishikawa, S., Steinberg, R. H., Naash, M. I., Duncan, J. L., Trautmann, N., … Flannery, J. G. (2018). Phenotypic characterization of P23H and S334ter rhodopsin transgenic rat models of inherited retinal degeneration. Exp Eye Res, 167, 56–90. doi:10.1016/j.exer.2017.10.023

Leinonen, H., Choi, E. H., Gardella, A., Kefalov, V. J., & Palczewski, K. (2019). A Mixture of U.S. Food and Drug Administration-Approved Monoaminergic Drugs Protects the Retina From Light Damage in Diverse Models of Night Blindness. Invest Ophthalmol Vis Sci, 60(5), 1442–1453. doi:10.1167/iovs.19-26560

Li, Z. Y., Kljavin, I. J., & Milam, A. H. (1995). Rod photoreceptor neurite sprouting in retinitis pigmentosa. J Neurosci, 15(8), 5429–5438.

Machida, S., Kondo, M., Jamison, J. A., Khan, N. W., Kononen, L. T., Sugawara, T., … Sieving, P. A. (2000). P23H rhodopsin transgenic rat: correlation of retinal function with histopathology. Invest Ophthalmol Vis Sci, 41(10), 3200–3209.

Marc, R. E., & Jones, B. W. (2003). Retinal remodeling in inherited photoreceptor degenerations. Mol Neurobiol, 28(2), 139–147. doi:10.1385/MN:28:2:139

Marc, R. E., Jones, B. W., Watt, C. B., & Strettoi, E. (2003). Neural remodeling in retinal degeneration. Prog Retin Eye Res, 22(5), 607–655.

Margolis, D. J., Newkirk, G., Euler, T., & Detwiler, P. B. (2008). Functional stability of retinal ganglion cells after degeneration-induced changes in synaptic input. J Neurosci, 28(25), 6526–6536. doi:10.1523/JNEUROSCI.1533-08.2008

Michalakis, S., Schaferhoff, K., Spiwoks-Becker, I., Zabouri, N., Koch, S., Koch, F., … Haverkamp, S. (2013). Characterization of neurite outgrowth and ectopic synaptogenesis in response to photoreceptor dysfunction. Cell Mol Life Sci, 70(10), 1831–1847. doi:10.1007/s00018-012-1230-z

Murphy, T. H., & Corbett, D. (2009). Plasticity during stroke recovery: from synapse to behaviour. Nat Rev Neurosci, 10(12), 861–872. doi:10.1038/nrn2735

Nikonov, S. S., Kholodenko, R., Lem, J., & Pugh, E. N., Jr. (2006). Physiological features of the S- and M-cone photoreceptors of wild-type mice from single-cell recordings. J Gen Physiol, 127(4), 359–374. doi:10.1085/jgp.200609490

Orban, T., Leinonen, H., Getter, T., Dong, Z., Sun, W., Gao, S., … Palczewski, K. (2018). A Combination of G Protein-Coupled Receptor Modulators Protects Photoreceptors from Degeneration. J Pharmacol Exp Ther, 364(2), 207–220. doi:10.1124/jpet.117.245167

Petersen-Jones, S. M. (1998). A review of research to elucidate the causes of the generalized progressive retinal atrophies. Vet J, 155(1), 5–18. doi:10.1016/s1090-0233(98)80028-2

Pfeiffer, R. L., Marc, R. E., & Jones, B. W. (2020). Persistent remodeling and neurodegeneration in late-stage retinal degeneration. Prog Retin Eye Res, 74, 100771. doi:10.1016/j.preteyeres.2019.07.004

Pozniak, P. D., White, M. K., & Khalili, K. (2014). TNF-alpha/NF-kappaB signaling in the CNS: possible connection to EPHB2. J Neuroimmune Pharmacol, 9(2), 133–141. doi:10.1007/s11481-013-9517-x

Prusky, G. T., Alam, N. M., Beekman, S., & Douglas, R. M. (2004). Rapid quantification of adult and developing mouse spatial vision using a virtual optomotor system. Invest Ophthalmol Vis Sci, 45(12), 4611–4616. doi:10.1167/iovs.04-0541

Ronning, K. E., Allina, G. P., Miller, E. B., Zawadzki, R. J., Pugh, E. N., Jr., Herrmann, R., & Burns, M. E. (2018). Loss of cone function without degeneration in a novel Gnat2 knock-out mouse. Exp Eye Res, 171, 111–118. doi:10.1016/j.exer.2018.02.024

Ross, J. W., Fernandez de Castro, J. P., Zhao, J., Samuel, M., Walters, E., Rios, C., … Kaplan, H. J. (2012). Generation of an inbred miniature pig model of retinitis pigmentosa. Invest Ophthalmol Vis Sci, 53(1), 501–507. doi:10.1167/iovs.11-8784

Sakami, S., Kolesnikov, A. V., Kefalov, V. J., & Palczewski, K. (2014). P23H opsin knock-in mice reveal a novel step in retinal rod disc morphogenesis. Hum Mol Genet, 23(7), 1723–1741. doi:10.1093/hmg/ddt561

Sakami, S., Maeda, T., Bereta, G., Okano, K., Golczak, M., Sumaroka, A., … Palczewski, K. (2011). Probing mechanisms of photoreceptor degeneration in a new mouse model of the common form of autosomal dominant retinitis pigmentosa due to P23H opsin mutations. J Biol Chem, 286(12), 10551–10567. doi:M110.209759 [pii] 10.1074/jbc.M110.209759

Sarria, I., Pahlberg, J., Cao, Y., Kolesnikov, A. V., Kefalov, V. J., Sampath, A. P., & Martemyanov, K. A. (2015). Sensitivity and kinetics of signal transmission at the first visual synapse differentially impact visually-guided behavior. Elife, 4, e06358. doi:10.7554/eLife.06358

Shen, N., Wang, B., Soto, F., & Kerschensteiner, D. (2020). Homeostatic Plasticity Shapes the Retinal Response to Photoreceptor Degeneration. Curr Biol. doi:10.1016/j.cub.2020.03.033

Stasheff, S. F. (2008). Emergence of sustained spontaneous hyperactivity and temporary preservation of OFF responses in ganglion cells of the retinal degeneration (rd1) mouse. J Neurophysiol, 99(3), 1408–1421. doi:10.1152/jn.00144.2007

Telias, M., Denlinger, B., Helft, Z., Thornton, C., Beckwith-Cohen, B., & Kramer, R. H. (2019). Retinoic Acid Induces Hyperactivity, and Blocking Its Receptor Unmasks Light Responses and Augments Vision in Retinal Degeneration. Neuron. doi:10.1016/j.neuron.2019.02.015

Tien, N. W., Soto, F., & Kerschensteiner, D. (2017). Homeostatic Plasticity Shapes Cell-Type-Specific Wiring in the Retina. Neuron, 94(3), 656–665 e654. doi:10.1016/j.neuron.2017.04.016

Toychiev, A. H., Ivanova, E., Yee, C. W., & Sagdullaev, B. T. (2013). Block of gap junctions eliminates aberrant activity and restores light responses during retinal degeneration. J Neurosci, 33(35), 13972–13977. doi:10.1523/JNEUROSCI.2399-13.2013

Tu, H. Y., Chen, Y. J., McQuiston, A. R., Chiao, C. C., & Chen, C. K. (2015). A Novel Retinal Oscillation Mechanism in an Autosomal Dominant Photoreceptor Degeneration Mouse Model. Front Cell Neurosci, 9, 513. doi:10.3389/fncel.2015.00513

Turrigiano, G. (2012). Homeostatic synaptic plasticity: local and global mechanisms for stabilizing neuronal function. Cold Spring Harb Perspect Biol, 4(1), a005736. doi:10.1101/cshperspect.a005736

Turrigiano, G. G. (2017). The dialectic of Hebb and homeostasis. Philos Trans R Soc Lond B Biol Sci, 372(1715). doi:10.1098/rstb.2016.0258

Turrigiano, G. G., Leslie, K. R., Desai, N. S., Rutherford, L. C., & Nelson, S. B. (1998). Activity-dependent scaling of quantal amplitude in neocortical neurons. Nature, 391(6670), 892–896. doi:10.1038/36103

Vinberg, F., Kolesnikov, A. V., & Kefalov, V. J. (2014). Ex vivo ERG analysis of photoreceptors using an in vivo ERG system. Vision Res, 101, 108–117. doi:10.1016/j.visres.2014.06.003

Vinberg, F., Wang, T., Molday, R. S., Chen, J., & Kefalov, V. J. (2015). A new mouse model for stationary night blindness with mutant Slc24a1 explains the pathophysiology of the associated human disease. Hum Mol Genet, 24(20), 5915–5929. doi:10.1093/hmg/ddv319

Wang, T., Pahlberg, J., Cafaro, J., Frederiksen, R., Cooper, A. J., Sampath, A. P., … Chen, J. (2019). Activation of rod input in a model of retinal degeneration reverses retinal remodeling and induces formation of functional synapses and recovery of visual signaling in the adult retina. J Neurosci. doi:10.1523/JNEUROSCI.2902-18.2019

Wang, T., Reingruber, J., Woodruff, M. L., Majumder, A., Camarena, A., Artemyev, N. O., … Chen, J. (2018). The PDE6 mutation in the rd10 retinal degeneration mouse model causes protein mislocalization and instability and promotes cell death through increased ion influx. J Biol Chem, 293(40), 15332–15346. doi:10.1074/jbc.RA118.004459

Young, R. W. (1985). Cell differentiation in the retina of the mouse. Anat Rec, 212(2), 199–205. doi:10.1002/ar.1092120215

