## Supplementary figures and images for "Homeostatic plasticity triggered by rod photoreceptor degenerative disease is associated with maintenance of sensitive night vision"

### Fig. 2. - Supplement 1. Activation of mitogen-activated protein kinase (MAPK) pathway in 1-mo P23H retinas. T-test: **P<0.01, ***P<0.001.

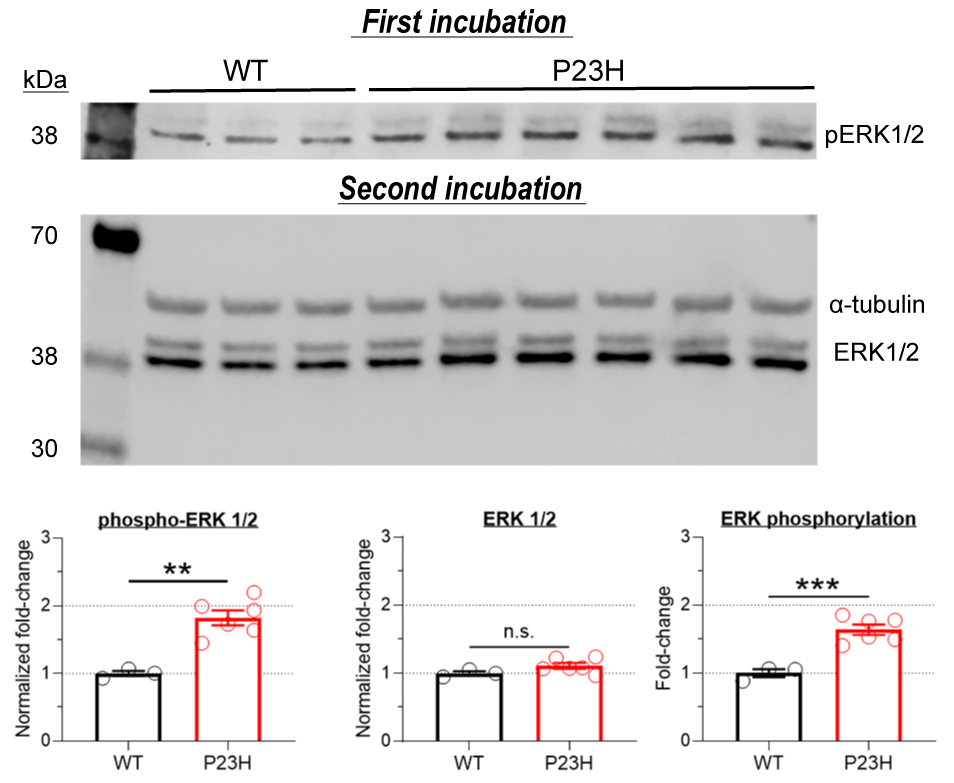
